# Tumor neoantigen heterogeneity thresholds provide a time window for combination immunotherapy

**DOI:** 10.1101/823856

**Authors:** Guim Aguadé-Gorgorió, Ricard Solé

## Abstract

Following the advent of immunotherapy as a possible cure for cancer, remarkable insight has been gained on the role of mutational load and neoantigens as key ingredients in T cell recognition of malignancies. However, not all highly mutational tumors react to immune therapies, and even initial success is often followed by eventual relapse. Recent research points out that high heterogeneity in the neoantigen landscape of a tumor might be key in understanding the failure of immune surveillance. In this work we present a mathematical framework able to describe how neoantigen distributions shape the immune response. Modeling T cell reactivity as a function of antigen dominancy and distribution across a tumor indicates the existence of a diversity threshold beyond which T cells fail at controling heterogeneous cancer growth. Furthemore, an analytical estimate for the evolution of average antigen clonality indicates rapid increases in epitope heterogeneity in early malignancy growth. In this scenario, we propose that therapies targeting the tumor prior to immunotherapy can reduce neoantigen heterogeneity, and postulate the existence of a time window, before tumor relapse due to de novo resistance, rendering immunotherapy more effective.

**Major Findings:** Genetic heterogeneity affects the immune response to an evolving tumor by shaping the neoantigen landscape of the cancer cells, and highly heterogeneous tumors seem to escape T cell recognition. Mathematical modeling predicts the existence of a well defined neoantigen diversity threshold, beyond which lymphocites are not able to counteract the growth of a population of highly heterogeneous subclones. Furthermore, evolutionary dynamics predict a fast decay of neoantigen clonality, rendering advanced tumors hard to attack at the time of immunotherapy. Within this mathematical framework we propose that targeted therapy forcing a selective pressure for resistance might as well increase neoantigen homogeneity, providing a novel possibility for combination therapy.

## I. INTRODUCTION

The mechanisms that make cancer a major cause of human death are deeply rooted in the principles governing the evolution of all life forms [1]. Through depletion of early mutations affecting mostly multicellular regulation and tissue homeostasis, cancer cells can overcome multiple selection barriers, making the human genome a pool for the evolution of a myriad possible phenotypes [2]. Highly diverse rogue populations within a tumor often include cells that resist, or evolve resistance to a vast array of therapy schemes [3,4]. Within this evolutionary paradigm, advanced, heterogeneous or metastatic cancer appears as a disease constantly evading a cure. Among other selective barriers, cancer cells often evolve the ability to evade immune surveillance [5], the otherways-normal capacity of the immune system to discern and attack non-self agents in the body [6] (Fig. 1a). Recent decades have seen a rapidly growing insight into the molecular details of immune silencing and tolerance [7], leading to the advent of checkpoint inhibition therapies targeting specific immune-evasion mechanisms in malignancies [8].

**FIG. 1:**
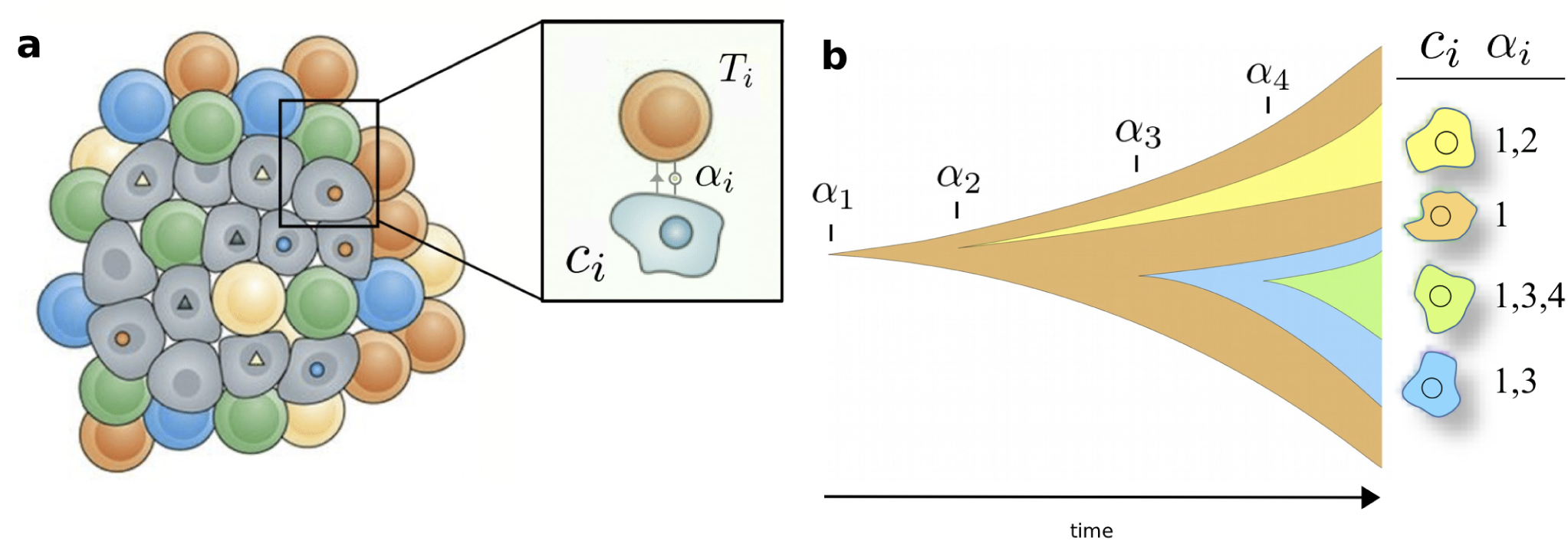
Neutral evolution of neoantigens after immune escape. Immune recognition and attack **(a)** effects a selective pressure towards activation of evasion mechanisms in cancer cells. After evasion, neoantigens (labeled *α_i_*) proliferate and evolve neutrally **(b)**. In our model, each color in **(b)** represents a subclone. After therapy reactivating the immune system, each subclone will die depending on its neoantigen composition; for example, the green subclone *c*_3_ harbors antigens 1, 3 and 4.

Once the immune system is back in place, the role of neoantigens, mutated surface proteins that elicit T cell recognition, has proven to be crucial, making genetic instability and mutational load valuable markers of eventual prognosis [9–12], and opening the door to novel therapy approaches targeting DNA stability to increase neoantigen production [13]. However, recent research highlights that not only the amount of neoantigens, but rather their heterogeneity and distribution across the tumor determines the eventual fate of immune therapies [14,15].

How do neoantigen distributions evolve within immune evaded tumors, and how does their shape affect the eventual outcome of T cell activation after therapy? Recent mathematical insight has emerged from the study of neutrality of neoantigen evolution as well as the effect of selective pressures towards immune escape [16]. Further research has proven it possible to characterize immunotherapy prognosis by quantitatively assessing the immunogeneicity of subclonal epitope composition [17]. However, how the degree neoantigen heterogeneity correlates with the immune response remains to be studied from a mathematical perspective.

In this work we introduce a mathematical framework of the cancer-immune interaction (Fig. 1a) that captures the effect of neoantigen evolution and heterogeneity on immune surveillance (Fig. 1b). The model predicts that neoantigen diversity shapes an all-or-none transition separating growing from immune-controled tumors. To understand how diversity arises during malignancy progression, we infer the evolutionary pace of average antigen clonality in a growing tumor, and eventually propose combination therapy approaches able to control the fast increase of tumor epitope heterogeneity.

## II. MATERIALS AND METHODS

### A. The role of neoantigen heterogeneity in the cancer-immune interaction

Mathematical models of immunology have been in place for decades [20]. In the field of cancer, relevant insight has been obtained from both differential equations and computational approaches, with particular descriptions of tumor dormancy or the role of inflammation, together with the existence of critical conditions separating cancer progression from its extinction (see [21] and references therein).

However, only recently have the models included genetic instability and neoantigen load as key drivers of immune recognition [22]. Recent approaches are also including neoantigen evolution [16] or subclonal structure and immunodominance effects [17] as parameters in the tumor-immune interaction. Can we make use of these results to understand the role of neoantigen clonality in immune control of malignancy?

Despite recent technological advances, tumor sequencing and neoantigen recognition algorithms cannot detect all antigens but only those present beyond a given frequency [23]. Within this picture, we describe the ecology of a tumor as a set of subclones characterized by harboring the same epitope composition, with subclonal-specific neoantigens over the detectability threshold together with those shared with its neighbouring phylogenetic branches or the whole tumor mass (Figure 1b). How does immune attack and cancer death correlate with neoantigen composition? From the set of neoantigens presented by a clone, only a very few are of significant immunogeneicity and elicit a strong T cell response [17], thus cancer cell death is expected to correlate with the most immunogenic antigen in their repertoire.

Previous research has already studied the neoantigenic capacity of epitopes, termed *neoantigen fitness* [17]. However, further work indicates that even highly recognizable antigens fail at inducing T cell response if they are not present in a sufficient fraction of the tumor [14]. Under this assumption, not only the recognition fitness of a neoantigen affects the subclone death, but also its frequency or distribution among the whole tumor (see Quick Guide to Model and Major Assumptions box for the complete model).

Prior modeling exercises have succesfully evaluated tumor evolution and response to immunotherapy under this perspective [17], but without explicitly considering heterogeneity in the antigen landscape. Furthermore, the process of evaluating the most dominant antigen in each clone, its recognition potential, and its distribution among the rest of subclones relies on bulk data processing where sampling biases due to localized biopsies [23] and imperfect antigen prediction [24] introduce a high level of noise. Several relevant questions emerge from these findings: Is it possible to gain insight into how T cells react at given neoantigen distributions without the need of neoantigen-specific computations? Are there common patterns stemming from our minimal model separating respondant from non-respondant patients, and what can we learn from them?

#### 1. Neoantigen Dominance D

To determine the immunogenic capacity of cancer neoantigens, a recent approach couples the likelihood of neoantigen presentation by the major histocompatibility complex (MHC) together with the recognition capacity by T cells, which stems from comparing epitope sequences with a database of known human antigens [17]. However, obtaining the whole neoantigen presentation map and recognition measures to evaluate expected subclone surveillance comes at a computational cost, and immunogeneicity of surface proteins is often overestimated [24]. Is there a simpler way to approximate the expected immunogeneicity, or recognition fitness, of a subclone considering their neoantigen load?

When studying the distribution of antigen fitness in (data from [17], antigen database collected from [9–11]), it can be seen that their distribution is highly skewed: a vast amount of neoantigens are of low, if not null, immunogeneicity, while a small set is highly immunodominant. We may ask, given the number of neoantigens present in a subclone, what is the probability that a highly immunodominant one has been generated? We build a process simulating the neutral production of neoantigens (see Box). From this, we obtain a first approximation of how average maximum fitness increases as more neoantigens are being produced.

Within this stochastic picture it is clear that, even in a scenario of a finite neoantigen landscape, many neoantigens have to be produced in order to find, on average, highly immunogenic ones. Simulations indicate that, for common subclones with about 1 to 200 antigens [17], recognition potential of the most dominant antigen scales linearly with the total neoantigen burden (Fig. 2a). For larger, more uncommon antigen loads, maximum immunogeneicity in simulations saturates, as the epitope database is finite.

**FIG. 2:**
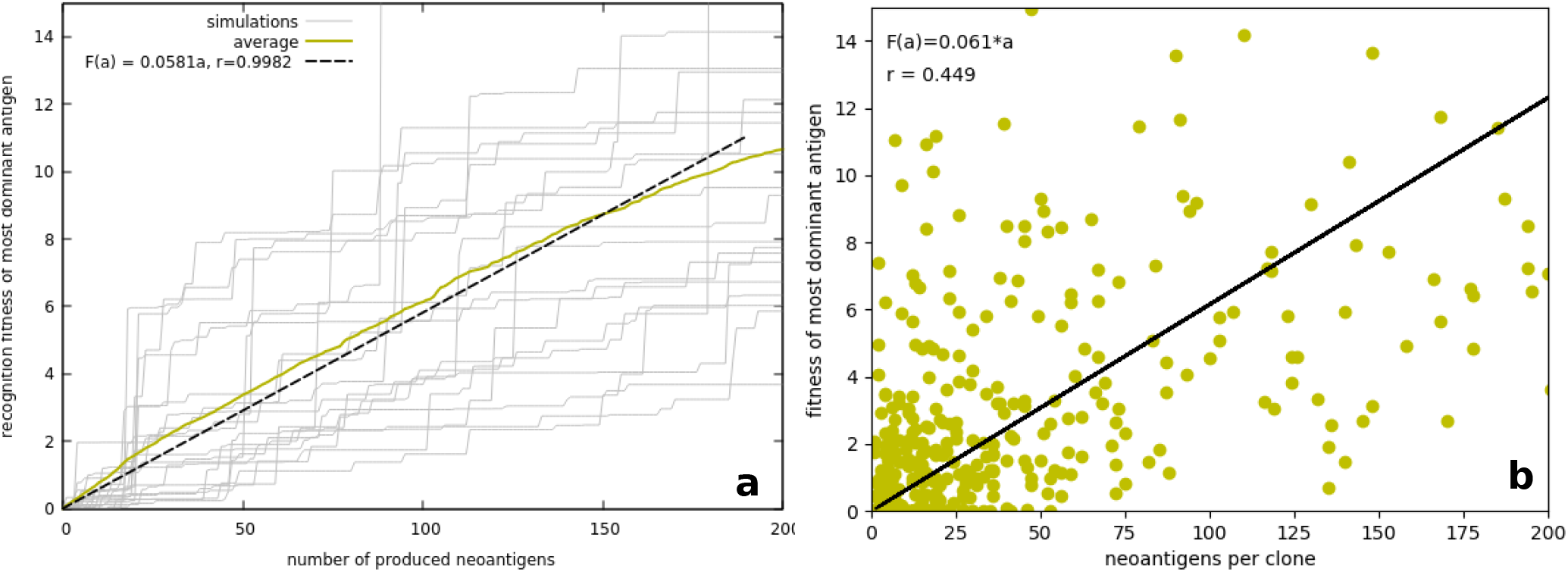
Correlation between subclonal neoantigen load and recognition potential of the fittest neoantigen. In **(a)**, simulations and simulation average show a linear increase as neoantigens are drawn and more dominant ones presented. In **(b)**, data from [17] is consistent with simulations. Both graphs are within the region of subclonal neoantigen load where most of the subclones are found (∼85%, data from [17]).

**FIG. 3:**
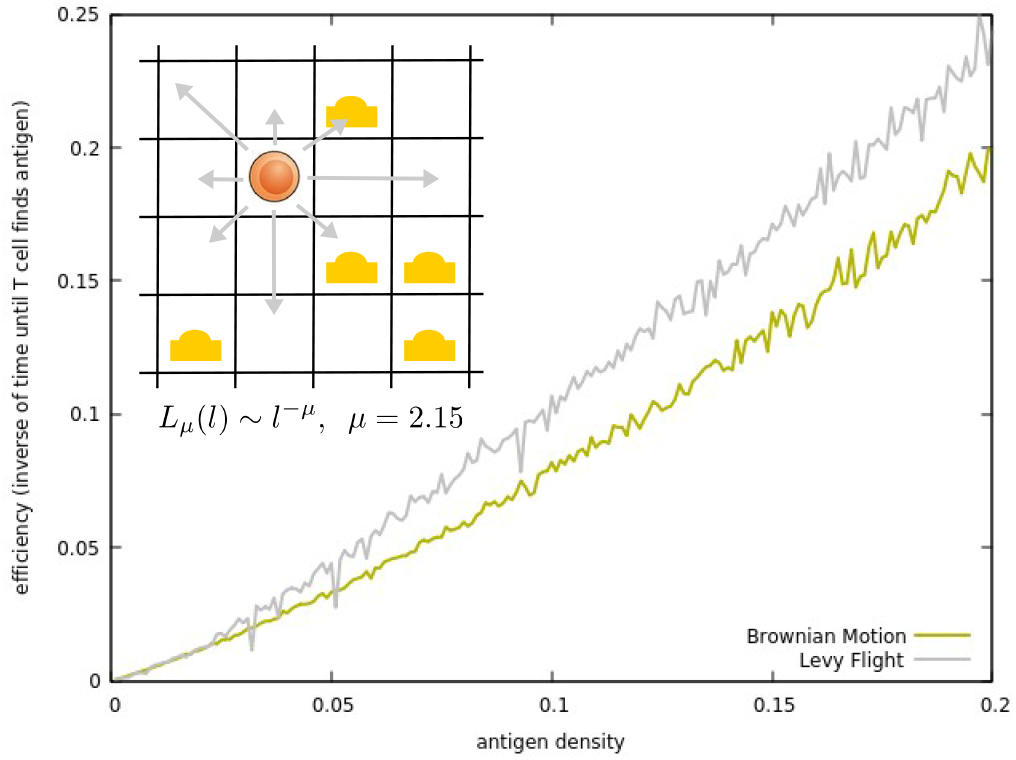
Spatial simulations for the effect of neoantigen frequency on T cell search efficiency. Neoantigens are distributed randomly on a two-dimensional grid according to their frequency. The inverse of the average time until a Lévy migrating T-cell search encounters the antigen scales linearly with the antigen frequency *γ_i_*. Simulations of Brownian-migrating T cells are presented for comparison, consistent with results indicating that Lévy strategies are more efficient for rare target search [18,27]

This approach can also be cross-tested by looking for positive correlations between the subclonal antigen burden and the dominant recognition potential of each subclone (Fig. 2b). The consistency between simulations and actual correlation supports the neutrality of neoantigen presentation [16,19]: if neoantigens where under strong selective pressures, one would expect deviations in Fig 2b from the simulations in Fig 2a indicating depletion of immunodominant antigens.

To summarize, this first minimal approach indicates the possibility of approximating a subclone immunogeneicity as a linear function of the antigen burden in that subclone

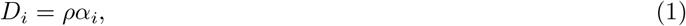

where *ρ*~0.06 from our simulations on random neoantigen presentation that are consistent with the correlation from the database in [17]. Despite the strong assumptions made, this approach gives us an initial way of estimating the possible immunogeneicity of a subclone without the need to assess neoantigen-specific *fitness*.

#### 2. Immune Search Efficiency E

Another key component in the immune response stems from the mechanisms of clonal selection. Briefly, upon presentation of a given antigen, helper T cells with the matching TCR replicate and release cytokines that eventually result in the expansion of a cytotoxic T cell clone [25]. This ensures efficient surveillance and further memory of previous antigenic encounters.

Within this scenario, the physical encounter of a cancer cell presenting given antigens and a corresponding T cell is the first necessary step to start the processes leading to eventual attack, and previous research has pointed out to modeling this search as a collision-rate process [26]. Following the relevance of neoantigen clonality pointed out in [14], we should ask if there is a relation between the concentration of given neoantigens in a tumor and the associated T cell dynamics. This can be approached by using ideas from the ecology of foraging [27]. In this context, it was shown that T cells could in fact migrate in search for pathogens following a generalized Lévy flight process [18]. These results seem to point towards lymphocites having evolved near-optimal search strategies for rare targets [18]. What are the implications on how the immune system faces an heterogeneous neoantigen tumor surface?

A spatial model for the cancer-immune interaction has been recently proposed in [28], where T cells walk on a two-dimensional grid in search for antigens. Here, we present a similar model able to explicitly compute the dependency of T cell search efficiency on antigen clonality. The specific construction of the stochastic simulations is described in the Box.

Despite efficiency obviously diverges as *γ*→1, a linear trend can be seen for small frequency levels similar to those found for the most common antigens in tumors [17]. Within our model, this translates into immune efficiency being a linear function of the clonality of the dominant antigen in the subclone, so that its death rate is linear with *γ_i_*. Together with neoantigen fitness, this gives a complete notion for the minimal dynamics of a cancer subclone

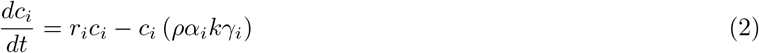

The factor *k* corresponds to the frequency-efficiency correlation *E* ~ *kγ* from our two-dimensional tumor surface simulations. Despite using real values from [18] for T cell motility, any quantitative estimation for *k* is beyond the scope of this work. As shown below, the linear relation between search efficiency and neoantigen distributions will suffice for our modeling results.

The model properly captures that subclones will die at rates corresponding to their number of antigens (as more antigens increase the probability of presenting a dominant one) and the frequency of their most dominant antigen across the tumor (T cell encounter increases with antigen clonality). Is this indicative of the role of neoantigen frequency on immune surveillance effectiveness?

### B. Neutral evolution of neoantigen distributions

The evolution of mechanisms to avoid T and B lymphocite recognition and destruction is nowadays recognized as a hallmark of cancer cells [6]. Early mutational processes generate surface proteins in malignant cells that activate immune surveillance, thus imposing a selective barrier overcomed by those cells able to avoid recognition [12,29]. What are the dynamics of neoantigen distributions once malignant cells avoid T cell attack?

Under the pressure of immune recognition, cancer cells with mechanisms to avoid lymphocite attack become selected, leading to tumors that are no longer controlled by the immune machinery [8]. Within this picture, the lack of an efficient immune response ensures that neoantigens are no longer negatively selected, making for tumors where neoantigen load and heterogeneity respond to neutral evolutionary dynamics.

The possibility of neutral evolution playing a role in cancer has seen remarkable progress [19]. In a recent work, Lakatos and colleagues develope a computational approach to understand the role of evolutionary dynamics on neoantigen distributions, and indicators of effectively-neutral evolution are found within neoantigen distributions in colorectal cancer exome sequencing data consistent with their simulations [16]. To gain insight in the dynamics of antigen distributions, we couple stochastic simulations, described in the Box, with the analytical estimates obtained in the Results section.

## III. RESULTS

### A. An heterogeneity threshold separates immune control from cancer outgrowth

We have seen how, with a minimal set of assumptions, we can obtain a dynamical description of how neoantigen load and particularly neoantigen frequency affect subclonal dynamics in the pressence of an effective immune response.Now further assumptions can be considered. On the one hand, assuming neoantigen neutrality, we can consider all clones to replicate at the same average rate, *r_i_* ~ *r* [16]. Additionally, the same neutrality argument can account for *γ_i_* ~ 〈*γ*〉: the frequency of dominant antigens at the time of immune attack should on average equal the average epitope frequency. To gain further insight into the complete tumor dynamics, we may ask if there are conditions under which all subclones can be controlled by immune surveillance, so that

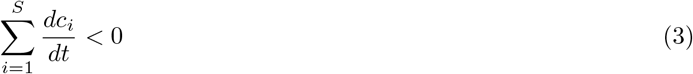

What is the actual upper limit *S* of the sum? As previously discussed, even if the distribution of neoantigens can account for epitopes only present in a single cell, realistic biopsies will hardly detect antigens with allelic frequency below certain cut-off levels specific of each sequencing method [23]. For those antigens with larger presence, we can separate the population into those *S* subclones with identical epitope configuration (Fig. 1b). This establishes the inequality:

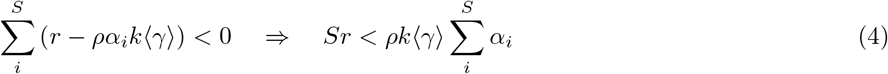

This result indicates the possibility of a threshold condition separating tumor growth from immune surveillance. Furthermore, this threshold results from basic tumor measures: average antigen frequency and average subclonal neoantigen load should outgrow the growth/recognition ratio

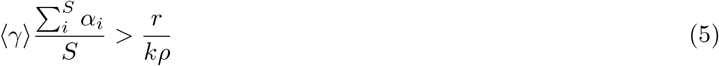

This result stemming from minimal considerations might have relevance in the light of recent research on immunotherapy prognosis. Despite not considering many features of the tumor composition, the existence a catastrophic shift separating tumor growth from extinction as a function of the average neoantigen clonality resembles recent clinical studies [14,15]. To which extent is our result comparable to experimental data?

By means of extracting specific neoantigen measures from the database in [17], we can study the correlations between months of survival in anti-PD-1-treated patients with lung cancer [9] and anti-CTLA-4 treated patients with melanoma [10,11] and several measures of neoantigen composition. Despite data being very sparse, it is plausible that the average antigen frequency times subclonal antigenic load of a tumor, as in our threshold, might render a better biomarker for the response to checkpoint inhibition therapy (Fig. 4), as compared with the nowadays accepted marker of total antigenic load.

**FIG. 4:**
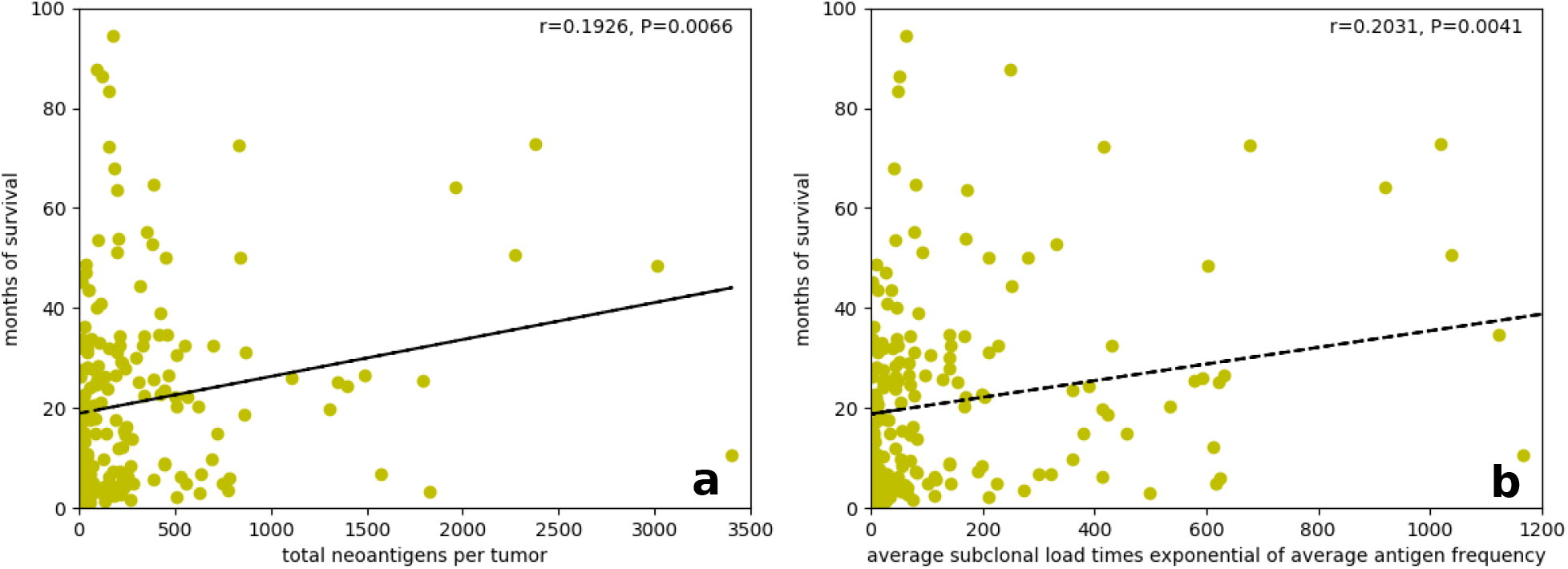
Correlation of patient biomarkers with months of survival after checkpoint inhibition therapy. In **(a)**, the common *total antigen load* marker. Including the average subclonal load and the average neoantigen frequency **(b)** seems to improve survival correlations, despite both data clouds being very sparse and P-value differences small (Data from [17]).

Knowing that neoantigen clonality shapes the immune recognition of a tumor, candidating for a possible prognosis biomarker, can we gain insight into how it evolves as the tumor grows prior to immune therapy? Is it possible to steer tumor evolution towards a more homogeneous neoantigen landscape?

### B. Average neoantigen frequency decays with tumor growth

Knowing that clonality might be a key aspect of T cell recognition, we use a mathematical modeling approach to understand the evolution of average antigen clonality in growing tumors. Following Gillespie simulations based on previous publications [16] as described in the Box, we study the analytical expressions able to characterize the evolution of antigen distributions.

In an exponentially growing tumor mass, we know that neoantigen load will obey (see [19])

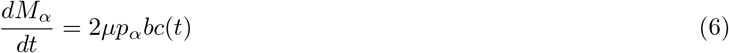

where *M_α_* is the amount of antigenic mutations in the tumor, *μ* stands for the overall mutational load and *p_α_* ~ 0.13 stands for the approximate portion of mutations predicted to bind MHC1 indicating possible immunogeneicity [25]. The 2 factor stems from the fact that, at each single birth event with rate *b*, both daughter cells in our simulations can undergo errors at rate *μp_α_*. Since the cancer population grows as *c*(*t*) = *c*_0_*exp*((*b − d*)*t*), we obtain

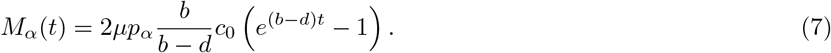

In the absence of cell death, the frequency of a given antigen will be directly related with the population size at presentation, *γ*(*α_i_*) = 1/*c*(*α_i_*) and remains constant [19]. When cells die out, this frequency can increase for antigens with surviving lineages [30] up to

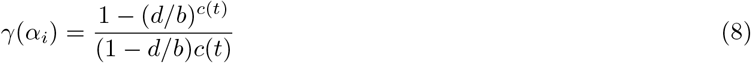

which is a relevant bias from 1/*c*(*t*) when birth and death rates are similar. Thus, in order to infer the frequency of each of the *M_α_* antigens, we need to know the tumor population when they were presented, and so their time of appearance. Each antigen appears when *M_α_* = *i* is a positive integer, which happens for

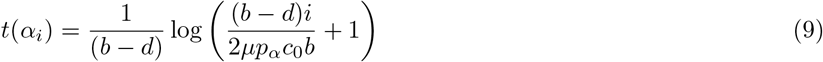

from where we infer that the population present when the *i*-th antigen is presented is

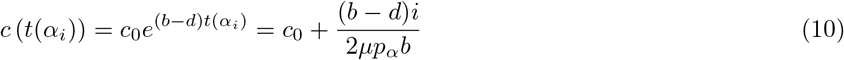

Both analytical estimates are consistent with simulations of exponentially growing tumors with random mutations (Fig. 5). However, an analytical sum of each antigen frequency (Eq. 9) using the population at appearance (Eq. 11) is not feasible. We still can, in the early tumor growth scenario where a blocked immune system does not jeopardize cell outgrowth, consider a low death-to-birth ratio and approximate the frequency of a mutation appearing at time *t_i_* by 1/*c*(*t_i_*) [19]. The average antigen clonality results from adding the inverse of each population at antigen appearance (Eq. 11) for all present antigens at time *t*. This results in a decaying average antigen frequency

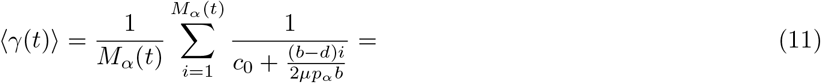

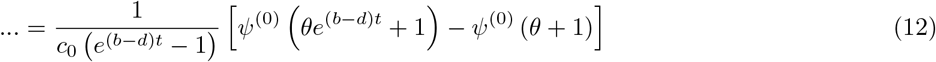

**FIG. 5:**
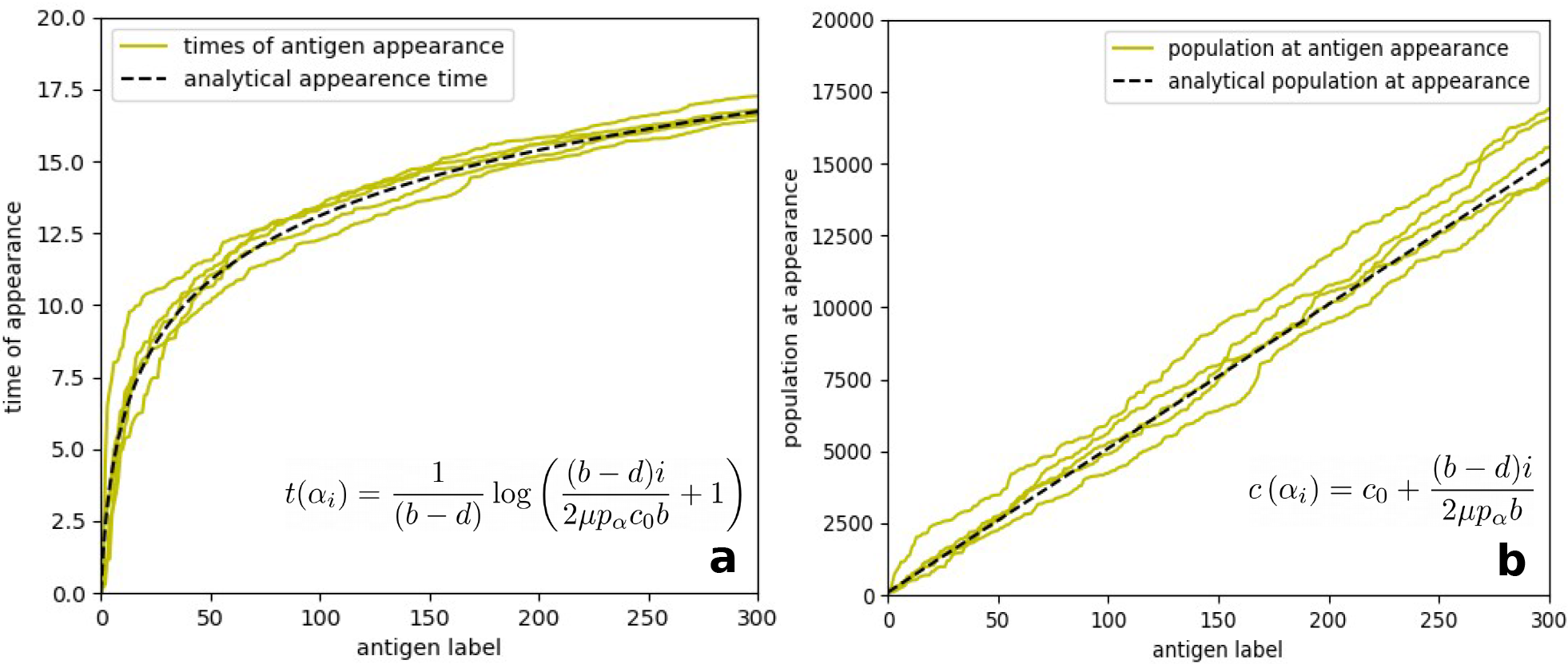
Simulations and analytical estimates for the average time **(a)** and total tumor population **(b)** at the appearance of each novel antigen presented in an exponentially growing tumor.

where *θ* = 2*μp_α_bc*_0_/(*b − d*) and *ψ*^(0)(*x*)^ is the digamma function [31].

This result can be compared to simulations (Fig. 6) and is a first analytical measure of the evolution of heterogeneity in the neoantigen landscape.

**FIG. 6:**
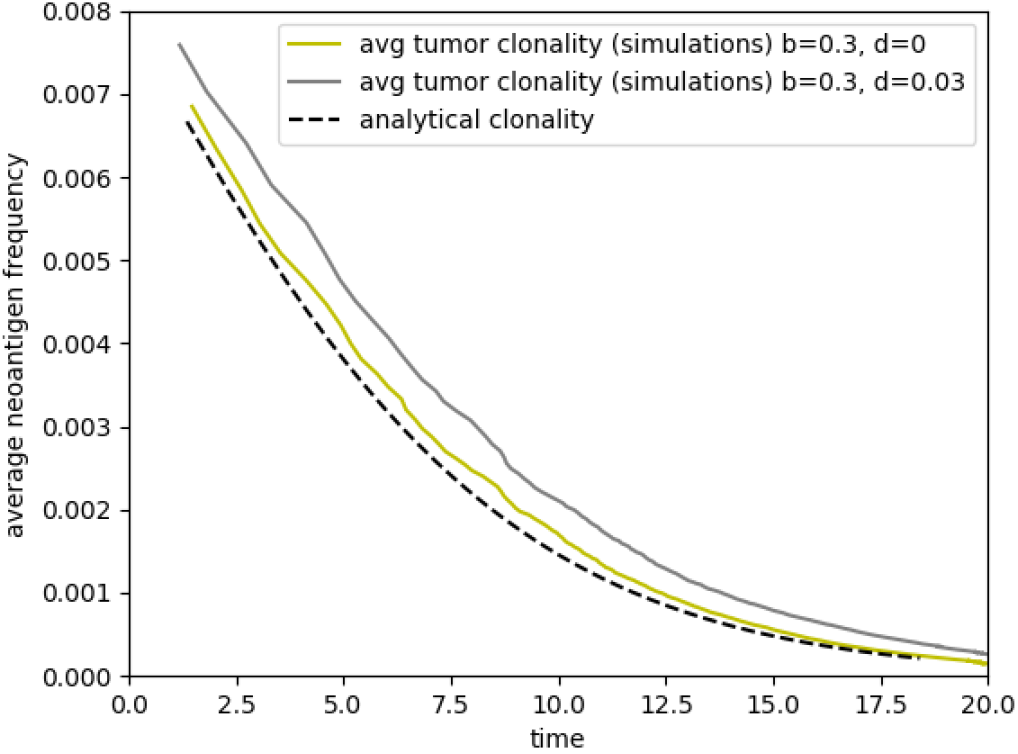
Evolution of average antigen frequency: analytical vs simulation results. In an exponentially growing tumor with *b* >> *d*, an analytical estimate for the decay of average clonality is found. This is a first estimate of the increase of heterogeneity in a neutraly evolving epitope landscape. Two runs, for *d* = 0 and *b/d* = 10 (as in [16]) are compared: a positive death rate increases the frequency of surviving antigens due to non-zero fixation probability [30], but the analytical result is still a good curvature estimate.

Knowing that average antigen clonality decays fast in growing tumors, and that the immune recognition threshold is sensible to such antigen clonality, it seems that this evolutionary dynamics might be related to the failure of immune activation in highly heterogeneous tumors. Is there any way in which neoantigen distributions can be steered into becoming more homogeneous prior to immune activation?

### C. Combination therapy could reduce neoantigen heterogeneity

Mathematical modeling indicates that heterogeneity thresholds exist, beyond which T cells might fail at reducing tumor growth, consistent with recent research [14,15]. Furthermore, our simulations and analytical estimates indicate that neoantigen clonality decreases rapidly during growth, meaning that checkpoint inhibition therapy activating the immune system will not necessarily succeed. Is there any way in which we can steer tumor evolution, so that neoantigen distributions become more homogeneous prior to immunotherapy? What is the possible effect of non-immune therapies on neoantigen frequency?

Consider now a general *in silico* therapy that is neutral to neoantigen load, as described in the Box and Figure 7d. As usual for many standard and targeted therapies, we include a set of mutations that render a subset of cells resistant to the therapy effect [32]. As therapy is administered, all cells without the mutation that ensures resistance will increase their death rate, thus leading to a general decay of tumor cell number (Figure 7a, 7d). This decrease in cell size comes with a halt in the production of novel antigens, while already present antigens are reduced as some may go extinct (Figure 7b).

**FIG. 7:**
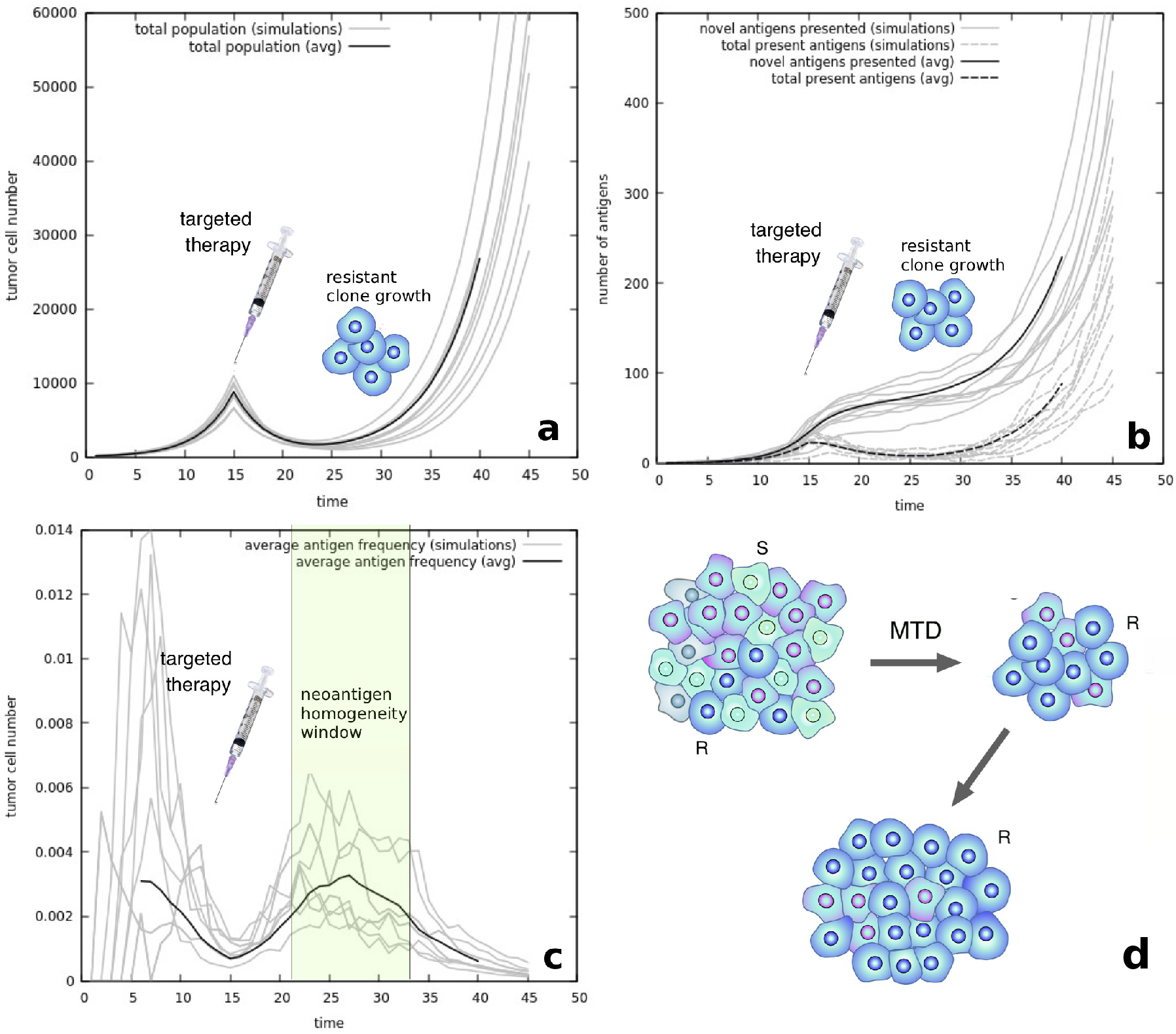
Evolutionary dynamics during non-immuno targeted therapy. **(a)**The population of cells decreases after therapy until a resistant subclone eventually repopulates the tumor. **(b)** Production of novel antigens slows down after therapy (black line) while total present antigens are reduced (dashed line). **(c)** Total tumor size decay produces an increase in clonality of those antigens present in the resistant subclone. Before the subclone repopulates the tumor, the increased average clonality creates a time window where the tumor is more neoantigen-homogeneous than prior to therapy, where checkpoint inhibition therapy might render more effective. **(d)** A scheme of the population dynamics shows subclonal outgrowth after maximum tolerated dose (MTD) treatment. We postulate that these process elicits an homogeneization of the neoantigen landscape.

In the light of these dynamics, what happens with average antigen clonality, key in surveillance effectiveness? It is interesting to see that, when therapy is administered, the overall reduction of cell number produces a rapid increase in average antigen frequency. Despite many antigens going extinct, those that belong to a surviving lineage (of cells that are resistant to therapy) increase their overall frequency due to death of non-resistant cells (Figure 7c). This result seems key in the effect of increased cell death due to neoantigen-neutral therapies.

As seen for many therapy approaches, a subpopulation of resistant cells often grows into a new, and often more malignant disease [3,4]. During this process, however, and until the resistant clone has grown into a full-size tumor, we observe a time window during which the average frequency of the neoantigen distribution is higher than when the cancer was of detectable size (Figure 7c). This evolutionary event indicates the possibility of a combination therapy where targeted therapy could be administered before checkpoint inhibitors, with the aim of reducing neoantigen heterogeneity and rendering the tumor more clonal and thus sensible to T cell attack.

## IV. DISCUSSION

The advent of immunotherapy approaches as a novel perspective for cancer cure has seen remarkable advances in recent years [33]. However, not all underlying mechanisms of immune surveillance and resistance are understood, and relapse seems to be present as cancer cells exploit evolutionary paths towards resistance [5,29]. A critical aspect of therapy efficiency lies in the role of neoantigen heterogeneity in immune surveillance [14,16,17]. How do neoantigen distributions evolve, and what is the effect of clonality in T cell response?

In the present work we have studied these problems through mathematical modeling. In particular, we have built a minimal model of heterogeneous cancer populations explicitly introducing their antigenic composition. Despite the large amount of ingredients governing adaptive immune recognition processes [34], we have described cell death by taking into account antigen dominancy and frequency. A minimal model indicates that dominancy could be a linear function of the total subclonal antigens, while the frequency of an antigen across a tumor might render a linear response on T cell activation for low clonalities.

Once the model is built, a diversity threshold separates cancer growth from immune control, and this threshold includes the measure of average antigen clonality, indicating the possibility for a novel biomarker for therapy prognosis and reinforcing recent experimental approaches [14,15]. Following the existence of a sharp transition governed by antigen frequency, we have studied how antigen frequency evolves prior to immunotherapy. Analytical estimates consistent with simulations indicate a fast decay of average antigen clonality owing to neutral dynamics. This might be indicative of why, at the time of checkpoint inhibition therapy, some cancers might not respond due to large heterogeneity levels in their antigen landscape [14,15].

An interesting point stemming from the model and simulations is that neoantigen distribution dynamics do not obey exponential or highly non-linear behaviours. If either immune search efficiency decayed exponentially with decreasing epitope frequency, or average clonality decayed exponentially during neutral antigen evolution, the model would predict that tumors rapidly escape from the region where T cell surveillance is still effective. Instead, smoother behaviours seem to govern the possible existence of an heterogeneity transition.

To overcome this threshold, we postulate the possibility of increasing epitope homogeneicity by selecting for a clone that is resistant to another therapy. Simulations indicate that, during the period where the rest of the tumor shrinks, positive selection for the resistant clone will ensure that average antigen homogeneity increases. Before the clone repopulates the tumor and further branching evolution arises, immunotherapy could render more effective as reactivated lymphocites would face a more homogeneous tumor.

Immunotherapy and its different combination approaches might be a tipping point in the search for a cure for cancer [35]. Complementing the advent of novel therapy methods, research on the mechanisms underlying resistance and relapse will prove crucial in the years to come [13,29]. The role of neoantigens and their heterogenous distributions across a tumor seems to be key in the response to checkpoint inhibition therapy. Multidisciplinary studies gaining insight into the dynamics of epitope heterogeneity and the possible existence of cooperative effects between targeted- and immuno-therapies might shed new light into increasing the effectiveness of immune surveillance across different tumor evolutionary histories and types [16,36].

## ACKNOWLEDGEMENTS

G.A. thanks Eszter Lakatos for interesting discussions on neoantigen evolution within the *Cancer Ecology and Evolution 2019* conference at the Wellcome Genome Campus. Both authors thank the Complex Systems Lab members as well as the HEF debates for insightful discussions. This work has been supported by the Botín-Foundation by Banco Santander through its Santander Universities Global Division, a MINECO grant FIS2015-67616-P (MINECO/FEDER, UE) fellowship, and AGAUR FI grant by the Universities and Research Secretariat of the Ministry of Business and Knowledge of the Generalitat de Catalunya and the European Social Fund and by the Santa Fe Institute.

## SUPPLEMENTARY GUIDE TO MODEL AND SIMULATIONS

### A. A minimal model for neoantigen-heterogeneous cancers

Our model stems from considering a metapopulation of cancer subclones, each described through a mean-field differential equation depending on the antigens they harbor (Fig. 1b)

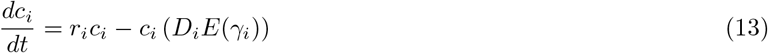

Our system describes the dynamics of each subclone, *c_i_*, composed by cells that replicate at rate *r_i_* and die at a rate depending on their subclonal antigen composition, which implicitly introduces the dynamics of the adaptive immune system without explicitly including it in the mean-field model as in [16,17]. This death rate will depend on the dominance *D* of the most dominant antigen in the subclone and the efficiency *E* of the immune system to detect such antigen, itself a function of the epitope frequency across the tumor bulk *γ*. Specific modeling is introduced to understand the shapes of both *D* and *E*.

#### 1. Neoantigen Dominance D

Recent methodologies have been able to estimate the capacity of individual neoantigens to elicit an immune response [17]. By coupling the likelihood of neoantigen presentation with the recognition capacity by T cells, a quantitative measure for each epitope is obtained. We study if there is a relation between recognition fitness of the most dominant antigen in a subclone and the total amount of antigens in it.

We build a stochastic simulation framework from a neoantigen database [17]. In the simulations, a cell adquires random mutations that select, one by one, antigens from the database. At each mutation, we update the recognition capacity to that of the novel epitope if it is the most dominant antigen at place to obtain a picture of how highly dominant antigens accumulate in a subclone (Figure 2).

#### 2. Immune Search Efficiency E

Experimental evidence suggests that the efficiency of the immune response depends on the distribution of neoantigens within the tumor. In principle, clonal antigens elicit a stronger response [14]. Following research on the mechanisms of lymphocite search [18], we build stochastic simulations to understand how the frequency of given epitopes in a tumor surface relates with the efficiency of T cell foraging. The model includes a two-dimensional grid where antigens are distributed, as a toy model for an immunogenic tumor surface. Antigens are characterized by the fraction of the cells they are present in, *γ_i_*. The grid is navigated by motile T cells displaying Lévy Flight migration statistics that will attack a cancer cell when a matching antigen is found. We compute how the relative presence of an antigen, *γ_i_*, relates with the necessary time until a *T − α_i_* encounter occurs. As a first approximation, we suppose that each T cell type responds only to one αi antigen, and map the efficiency of the search (the inverse of average time until first T-*α_i_* encounter) as a function of the antigen density *γ_i_* (Figure 3).

### B. Neoantigen evolution after immune evasion and targeted therapy

Neutral evolution seems to govern neoantigen dynamics after immune evasion mechanisms are selected for [16, 19]. A stochastic model for the evolution of neoantigens in a growing tumor is built to verify our analytical estimates (Fig. 1b). Similar to that in [16], our model consists in a birth-death Gillespie simulation of exponential growth of cancer cells, that replicate at rate *b* and die at rate *d*. At each replication event, both daughter cells can gain a set of mutations taken from a Poisson distribution with parameter *μ*. From this set of mutations, *p_α_* are considered to be of immunogenic capacity and produce a neoantigen. In our simulations, each antigen is associated to the mutated cell and labeled with a natural number *α*_1_, *α*_2_, *α*_3_, *α*_4_… (Fig. 1b). We record the population size, number of generated and present antigens and the frequency of each in the population. This last measure can be used to obtain the evolution of average antigen frequency along tumor growth. To understand the effect of therapy in neoantigen distributions, we introduce a type of mutations that confer resistance to a given drug. Within the space of possible mutations, some are considered to confer resistance with probability *p_R_*. Once therapy is administered, sensitive cells will die at rate *d_S_*, larger than their natural rate *d*, while resistant cells will die out at rate *d_R_*, much smaller. The introduction of this novel dynamics results in the reduction of tumor size followed by the growth of a resistant, selected subclone. We follow the dynamics of antigen number and frequency during this process to understand the effects of targeted therapy and selection for resistance in tumor growth.

## Notes

The authors declare no potential conflicts of interest.

## REFERENCES

1. Greaves M, Maley CC. Clonal evolution in cancer, Nature 2012;481:306–13

2. Negrini S, Gorgoulis VG, Halazonetis TD. Genomic instability–an evolving hallmark of cancer. Nat Rev Mol Cell Biol 2010;11:220–8.

3. Holohan C, Schaeybroeck SV, Daniel B, Longley DB, Johnston PG. Cancer drug resistance: an evolving paradigm. Nat Rev Cancer 2013;13:714–26

4. Burell RA, Swanton C. Tumour heterogeneity and the evolution of polyclonal drug resistance. Mol Oncol 2014;8(6):1095–111.

5. Ribas A. Adaptive Immune Resistance: How Cancer Protects from Immune Attack Cancer Discov 2015;5(9):915–19.

6. Hanahan D, Weinberg RA. Hallmarks of cancer: The next generation. Cell 2011;144(5):646–74.

7. Xing Y, Hongquist KA. T-cell tolerance: central and peripheral. Cold Spring Harb Perspect Biol 2012;4:6.

8. Ribas A, Wolchok JD. Cancer immunotherapy using checkpoint blockade. Science 2018;359(6382):1350–55.

9. Rizvi NA, Hellmann MD, Snyder A, Kvistborg P, Makarov V, Havel JJ, et al. Mutational landscape determines sensitivity to PD-1 blockade in nonsmall cell lung cancer. Science 2015;348:124–8.

10. Snyder A, Makarov V, Merghoub T, Yuan J, Zaretsky JM, Desrichard A, et al. Genetic basis for clinical response to CTLA-4 blockade in melanoma. N Engl J Med 2014;371:2189–99.

11. Van Allen EM, Miao D, Schilling B, Shukla SA, Blank C, Zimmer L, et al. Genomic correlates of response to CTLA4 blockade in metastatic melanoma. Science 2015;aad0095.

12. Schumacher TN, Schreiber RD. Neoantigens in cancer immunotherapy. Science 2015;348:69–74.

13. Zappasodi R, Merghoub T, Wolchok JD. Emerging Concepts for Immune Checkpoint Blockade-Based Combination Therapies. Cancer Cell 2018;33(4):581–98.

14. McGranahan N, Furness A, Rosenthal R, Ramskov S, Lyngaa R, Saini SK, et al. Clonal neoantigens elicit T cell immunoreactivity and sensitivity to immune checkpoint blockade. Science 2016;351:1463–9.

15. Wolf Y, Bartok O, Patkar S, Eli GB, Cohen S, Litchfield K, et al. UVB-Induced Tumor Heterogeneity Diminishes Immune Response in Melanoma. Cell 2019;179(1):219–35.

16. Luksza M, Riaz N, Makarov V, Balachandran VP, Hellmann MD, Solovyov A, et al. A neoantigen fitness model predicts tumour response to checkpoint blockade immunotherapy. Nature 2017;551:517–20.

17. Lakatos E, Williams MJ, Schenck RO, Cross WCH, Househam J, Werner B, et al. Evolutionary dynamics of neoantigens in growing tumours. BioRxiv 209;536433: https://doi.org/10.1101/536433

18. Harris TH, Banigan EJ, Christian DA, Konradt C, Tait Wojno ED, Norose K, et al. Generalized Lvy walks and the role of chemokines in migration of effector CD8+ T cells. Nature 2012; 486(7404):545–48.

19. Williams MJ, Werner B, Barnes CP, Graham TA, Sottoriva A. Identification of neutral tumor evolution across cancer types. Nat Gen 2016;48:238–44.

20. Perelson AS, Weisbuch G. Immunology for physicists. Rev Mod Phys 1997;69:1219.

21. Eftimie R, Bramson JL, Earn DJD. Interactions Between the Immune System and Cancer: A Brief Review of Non-spatial Mathematical Models. Bull Math Biol 2011;73:2–32.

22. Aguadé-Gorgorió G, Solé RV. Genetic instability as a driver for immune surveillance. bioRxiv 2019;527689: https://doi.org/10.1101/527689

23. Stead LF, Sutton KM, Taylor GR, Quirke P, Rabitts P. Accurately Identifying LowAllelic Fraction Variants in Single Samples with NextGeneration Sequencing: Applications in Tumor Subclone Resolution. Hum Mutat 2013;34:1432–8.

24. Yadav M, Jhunjhunwala S, Phung QT, Lupardus P, Tanguay J, Bumbaca S, Franci C, et al. Predicting immunogenic tumour mutations by combining mass spectrometry and exome sequencing. Nature 2014;515:572–6.

25. Starr TK, Jameson SC, Hogquist KA. Positive and Negative Selection of T Cells. Annu Rev Immunol 2003;21:139–176.

26. Budhu S, Loike JD, Pandolfi A, Han S, Catalano G, Constantinescu A, et al. CD8+ T cell concentration determines their efficiency in killing cognate antigenexpressing syngeneic mammalian cells in vitro and in mouse tissues. J Exp Med 2010;207(1):223–35.

27. Viswanathan GM, Buldyrev SV, Havlin S, da Luz MGE, Raposo EP, Stanley HE. Optimizing the success of random searches. Nature 1999;401:911–14.

28. Macfarlane FR, Chaplain MA, Lorenzi T. A stochastic individual-based model to explore the role of spatial interactions and antigen recognition in the immune response against solid tumours. J Theor Biol 2019;480:43–55.

29. Sharma P, Hu-Lieskovan S, Wargo JA, Ribas A. Primary, adaptive, and acquired resistance to cancer immunotherapy. Cell 2017;168:707–23.

30. Bozic I, Gerold JM, Nowak MA. Quantifying Clonal and Subclonal Passenger Mutations in Cancer Evolution. PLOS Comp Biol 2016;12(2):1–19.

31. Abramowitz M, Stegun IA (eds). “6.3 psi (Digamma) Function.” Handbook of Mathematical Functions with Formulas, Graphs, and Mathematical Tables. New York: Dover 1972:258–9.

32. Sawyers C. Targeted cancer therapy. Nature 2004;432:294–97.

33. Rosenberg SA. Entering the mainstream of cancer treatment. Nat Rev Clin Oncol 2014;11(11):630–2.

34. Gajewski TF, Schreiber H, Fu YX. Innate and adaptive immune cells in the tumor microenvironment Nat Immunol 2013;14:1014–22.

35. Hodge JW, Ardiani A, Farsaci B, Kwilas AR, Gameiro SR. The Tipping Point for Combination Therapy: Cancer Vaccines With Radiation, Chemotherapy, or Targeted Small Molecule Inhibitors. Sem Oncol 2012;39(3):323–339.

36. Park DS, Robertson-Tessi M, Luddy KA, Maini PK, Bonsall MB, Gatenby RA et al. The Goldilocks Window of Personalized Chemotherapy: Getting the Immune Response Just Right. Cancer Res 2019;79(20):5302–15.

